# *Arabidopsis* bZIP11 is a susceptibility factor during *Pseudomonas syringae* infection

**DOI:** 10.1101/2020.11.25.398727

**Authors:** Matthew J. Prior, Jebasingh Selvanayagam, Jung-Gun Kim, Monika Tomar, Martin Jonikas, Mary Beth Mudgett, Sjef Smeekens, Johannes Hanson, Wolf B. Frommer

**Affiliations:** Department of Botany and Plant Sciences, University of California Riverside, Riverside, CA 92507, USA; Department of Plant Biology, Carnegie Institution for Science, Stanford, CA 94305, USA; Department of Biology, Stanford University, Stanford, CA 94305, USA; Molecular Plant Physiology, Department of Biology, Utrecht University, Utrecht, The Netherlands; Department of Molecular Biology, Princeton University, 119 Lewis Thomas Laboratory, Washington Road, Princeton, NJ, USA; Umeå Plant Science Centre, Department of Plant Physiology, Umeå University, Umeå, Sweden; New address: Molecular Physiology, Heinrich Heine Universität, Düsseldorf, and Max Planck Institute for Plant Breeding Research, Köln, Germany

**Keywords:** Bacterial pathogenesis, susceptibility factor, bZIP transcription factor, plant nutrient secretion systems

## Abstract

The induction of plant nutrient secretion systems is critical for successful pathogen infection. Some bacterial pathogens, *e.g. Xanthomonas* species, use TAL (transcription activator-like) effectors to induce transcription of SWEET sucrose efflux transporters. *Pseudomonas syringae* pathovar (pv.) *tomato* strain DC3000 lacks TAL effectors, yet is able to induce multiple SWEETs in *Arabidopsis thaliana* by unknown mechanisms. Since bacteria require other nutrients besides sugars for efficient reproduction, we hypothesized that *Pseudomonas* may depend on host transcription factors involved in secretory programs to increase access to essential nutrients. Bioinformatic analyses identified the *Arabidopsis* basic-leucine zipper transcription factor bZIP11 as a potential regulator of nutrient transporters, including SWEETs and UmamiT amino acid transporters. Inducible downregulation of *bZIP11* expression in *Arabidopsis* resulted in reduced growth of *P. syringae* pv. *tomato* strain DC3000, whereas inducible overexpression of *bZIP11* resulted in increased bacterial growth, supporting the hypothesis that bZIP11 regulated transcription programs are essential for maximal pathogen titer in leaves. Our data are consistent with a model in which a pathogen alters host transcription factor expression upstream of secretory transcription networks to promote nutrient efflux from host cells.

## Introduction

Plant pathogens face two distinct challenges with respect to successful reproduction during invasion of their hosts. First, they must evade or suppress the host immune response to survive. Second, they must rapidly acquire sufficient nutrients to fuel their growth, often to very high titers in the host tissue. Substantial efforts provide a detailed picture of how plant pathogens overcome immune responses (Dou and Zhou 2012; Jones and Dangl 2006); however, we are only beginning to understand the strategies pathogens employ to secure access to nutrients from their hosts (Bezrutcyk et al, 2017).

One strategy used by bacterial pathogens of the genus *Xanthomonas* to access plant nutrients is the deployment of TAL (transcription activator-like) effectors. TALs induce the transcription of SWEET sucrose transporters in diverse hosts (*e.g.* rice, cassava, cotton and pepper), where each TAL effector induces a specific host sugar transporter gene (Chen et al. 2010; Cohn et al. 2014; Cernadas et al. 2014; Cox et al. 2017). TAL-induced expression of rice *OsSWEET* genes is required for *Xanthomonas oryzae* pv. *oryzae* (*Xoo*) pathogenesis, indicating that SWEETs are key host susceptibility factors during *Xanthomonas-*rice interactions (Chen et al. 2010; Cernadas et al. 2014). Rice lines carrying mutations in the TAL effector-binding site in the promoters of *OsSWEET11, 13* or *14* are resistant to *Xoo* (Chen et al. 2010; Cernadas et al. 2014). Promoter variants in *SWEET* genes yield broad spectrum resistance against bacterial leaf blight (Eom et al. 2019; Oliva et al. 2019). *SWEET*s are also induced in other plants in response to diverse pathogens, including bacteria, protists and fungi (Chen et al. 2010; Siemens et al. 2006; Chong et al. 2014; Cox et al. 2017). The genomes of these pathogens apparently lack TAL effector homologs (Buell et al. 2003; Schwelm et al. 2015; Hahn et al. 2014), implicating alternate mechanisms for the induction of *SWEET* genes.

Given that pathogens require access to essential micro- and macroelements in addition to sugars for growth, we hypothesized that pathogens may target transcription factors that alter nutrient availability at the local site of infection. To test this hypothesis, we asked whether the apoplasmic pathogen *Pseudomonas syringae* pv. *tomato* strain DC3000 induces multiple nutrient transporters, including SWEETs, in an effector-dependent manner (*i.e.* requiring the use of its Type Three Secretion System (T3SS)). We used a bioinformatic approach, reasoning that *P. syringae* pv. *tomato* strain DC3000 might upregulate the expression of multiple *SWEET*s and/or other nutrient transporters by targeting a transcription factor that normally functions upstream of metabolic programs found in secretory cell types such as seed coat or tapetum. We predicted that ectopic induction of such a transcription factor in local leaf cells could effectively increase the metabolic pools of nutrients in the reprogrammed cells and simultaneously induce transporters to efflux nutrients into the apoplasm for extracellular pathogens like *P. syringae* pv. *tomato* strain DC3000 (Fig. 1A).

**Figure 1.**
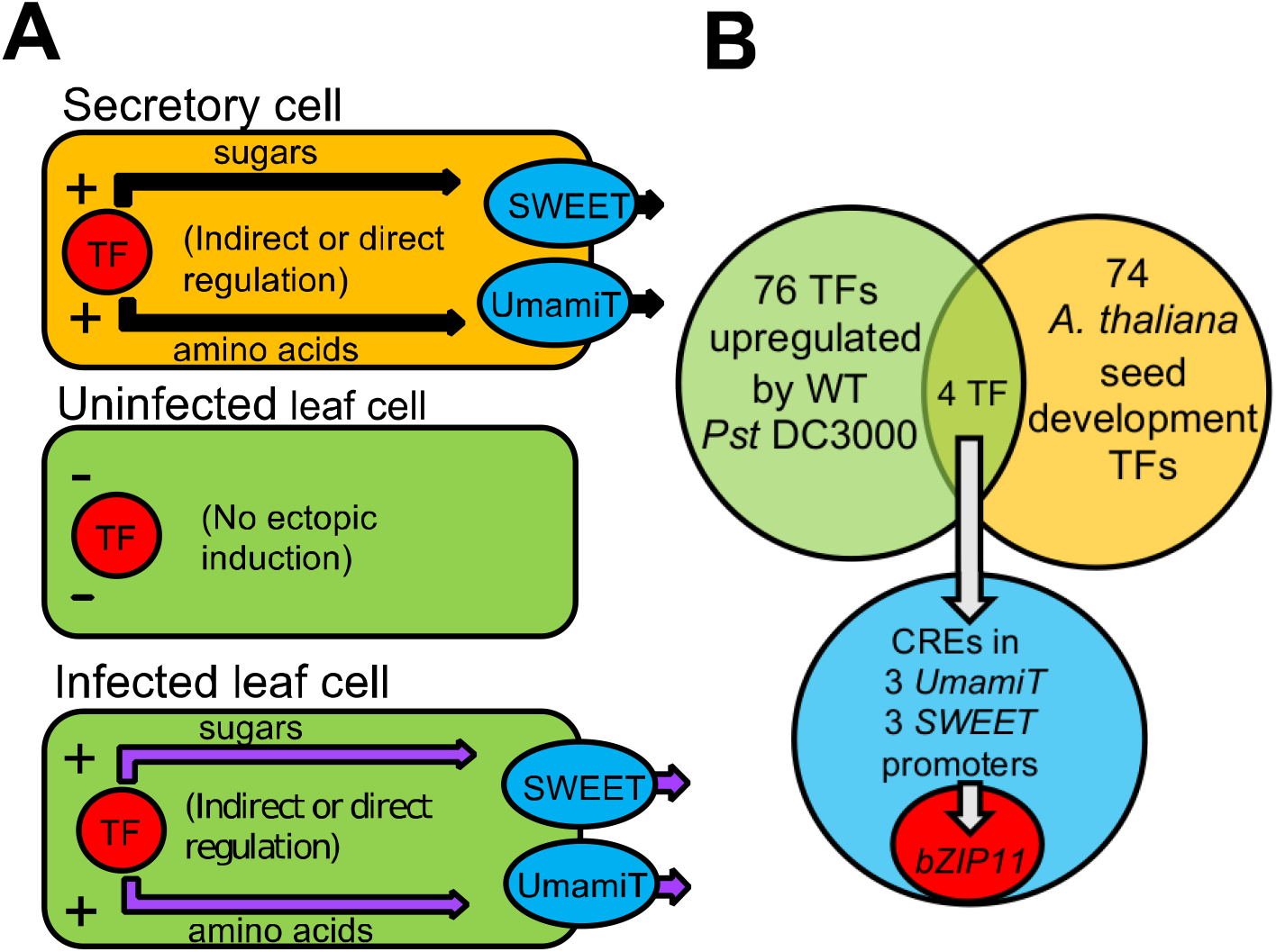
Screen for a transcription factor (TF) that influences secretory cell functions and is induced by the bacterial pathogen *P. syringae* pv. *tomato* strain DC3000. *(A)* Diagram showing basic movement of nutrients from secretory cells during seed development (top), and those proposed in uninfected leaf cells (middle) and infected leaf cells (bottom) by a TF that influences downstream *SWEET* and *UmamiT* transcription. *(B)* Bioinformatic analysis of microarray data compared 74 TFs previously identified in distinct groups by hierarchical clustering during *Arabidopsis* seed development against 76 TFs induced by *Pst* DC3000 in a T3SS-dependent manner in *Arabidopsis* leaves. One of the four TFs found in both datasets, *bZIP11* was identified as a single best candidate TF.

*Arabidopsis thaliana* bZIP11, a basic leucine-zipper (bZIP) domain transcription factor is known to be a crucial transcriptional regulator of genes involved in plant sugar and amino acid metabolism (Hanson et al. 2008; Ma et al. 2011). Here, we report that bZIP11 acts as a susceptibility factor during *P. syringae* pv. *tomato* strain DC3000 infection. We show that during development bZIP11 influences the transcription of multiple nutrient transporters with hallmarks of nutrient efflux from secretory cells. We also show that *bZIP11* expression is required for maximal *P. syringae* pv. *tomato* strain DC3000 growth in *Arabidopsis* leaves and that overexpression of bZIP11 leads to increased bacterial growth. Our data suggest that bZIP11 regulated transcription programs play an important in bacterial pathogenesis.

## Results

### Bioinformatic analyses identified *bZIP11* mRNA induction during infection

Three criteria were used to bioinformatically identify transcription factors in *Arabidopsis* that are selectively induced by pathogens, and which in turn could ectopically induce transcription of transporter genes for nutrient efflux, either directly or indirectly (Fig. 1A). First, we examined a set of transcription factors expressed in distinct hierarchical clusters during *Arabidopsis* seed development, as the seed coat is a secretory cell layer that expresses specific nutrient exporters, seen in Table S9 of the study by Le et al. 2010, and in particular SWEET sucrose and UmamiT amino acid transporters (Besnard et al. 2018; Chen et al. 2015; Müller et al. 2005). We then compared this group to another set of transcription factors reported to be induced in *Arabidopsis* leaves during *P. syringae* pv. *tomato* strain DC3000 in T3SS-dependent manner, found in Table 3 of the study by Truman et al. 2006. The overlap between these two sets identified four candidate transcription factors: bZIP11, bZIP9, AGL18, and At3g19910 (Fig. 1B, Table S1). We identified transcription factor binding sites (*i.e. cis* regulatory elements or CREs) enriched in promoters of putative nutrient transporters including three *SWEET*s and three *UmamiT*s and found an enrichment of bZIP sites (Table S2) (Zambelli et al. 2009). Of these six transporters, SWEET15, UmamiT18, and UmamiT29 have known feeding roles during seed development, raising the possibility that they might also play a role during pathogen infection (Chen et al. 2015; Ladwig et al. 2010; Müller et al. 2005).

Among the candidates identified, we selected bZIP11 as the top candidate for further studies because it fulfilled all of our bioinformatic and physiological criteria: (i) expressed in a secretory tissue (such as seed coat during seed filling), (ii) induced by the pathogen, (iii) and the candidate gene list is enriched for bZIP binding sites in their promoters. We also prioritized bZIP11 because its expression is regulated translationally by sucrose levels and is known to be crucial for controlling the carbon and nitrogen balance and regulating amino acid metabolism (Hanson et al. 2008; Ma et al. 2011). Prior work showed that overexpression of bZIP11 in *Arabidopsis* resulted in a marked increase in metabolic pools towards substrates for glycolysis, including increased glucose, fructose, and sucrose content, while multiple amino acids concentrations are also increased (Ma et al. 2011). Moreover, overexpression of a tobacco *bZIP11* homolog resulted in increased leaf sugar levels (Thalor et al. 2012). Taken together, the phenotypes observed in plants with bZIP up- and downregulation are consistent with functional characteristics of a transcription factor that alters metabolite levels and transporter gene expression.

### bZIP11 is induced during *P. syringae* pv.*tomato* strain DC3000 infection in an effector dependent manner

An analysis of published microarray datasets indicated that in three of four studies, *bZIP11* was induced by *P. syringae* pv. *tomato* strain DC3000 in a T3SS effector dependent manner (Truman et al. 2006; Thilmony et al. 2006; Kemmerling et al. 2007; Cumbie et al. 2011; Huang et al. 1992). To independently validate these findings, real time PCR analyses were performed using *Arabidopsis* Col-0 leaves infected with wild-type *P. syringae* pv. *tomato* strain DC3000 or *P. syringae* pv. *tomato* strain DC3000 Δ*hrcU*, which is defective in Type Three secretion (Huang et al. 1992). In three separate biological repeats, *bZIP11* mRNA levels were significantly increased in leaves following inoculation with wild-type *P. syringae* pv. *tomato* strain DC3000 but not the *P. syringae* pv. *tomato* strain DC3000 Δ*hrcU* mutant, indicating T3SS-dependence (Supplementary Fig. S1). Taken together, we hypothesize that *P. syringae* pv. *tomato* strain DC3000 uses T3SS effector proteins to induce *bZIP11* transcription.

### bZIP11 is necessary for normal development, metabolism, and transporter gene expression

To test the importance of bZIP11 in pathogen susceptibility, estradiol inducible amiRNA *Arabidopsis* lines targeting *bZIP11* were generated, as homozygous null mutants could not be recovered. Two independent transgenic lines (*bzip11*-s1, *bzip11*-s2) were chosen after analyzing *bZIP11* mRNA abundance by real time PCR (Supplementary Fig. S2A). With the addition of 5 μM estradiol, mRNA abundance in *bzip11*-s1 and *bzip11*-s2 was approximately 69% and 42% of wild-type levels, respectively (Supplementary Fig. S2A). Both amiRNA lines were smaller in stature compared to the wild-type Col-0 background at 4 weeks of growth in soil, consistent with expression of the amiRNA without the hormone (Supplementary Fig S2B). This could be due to a positional effect of insertion of the transgene into the chromosome or could be due to a low level of constitutive repression of *bZIP11* in absence of estradiol (Supplementary Fig. S2B). When germinated on 50 μM estradiol-containing media, both *bzip11*-s1 and *bzip11*-s2 seedlings showed chlorotic leaves and compromised root and shoot growth compared to wild-type Col-0 (Fig. 2A, 2B). Addition of sucrose to media containing 50 μM estradiol rescued the growth defects of the *bzip11*-s1 amiRNA line (Fig. 2E). These results intimate that bZIP11 is essential for normal growth of *Arabidopsis* and its effects can be bypassed through the addition of carbohydrates. They also are consistent with a role of bZIP11 upstream of the sucrose synthesis and/or export during growth regulation in *Arabidopsis*.

**Figure 2.**
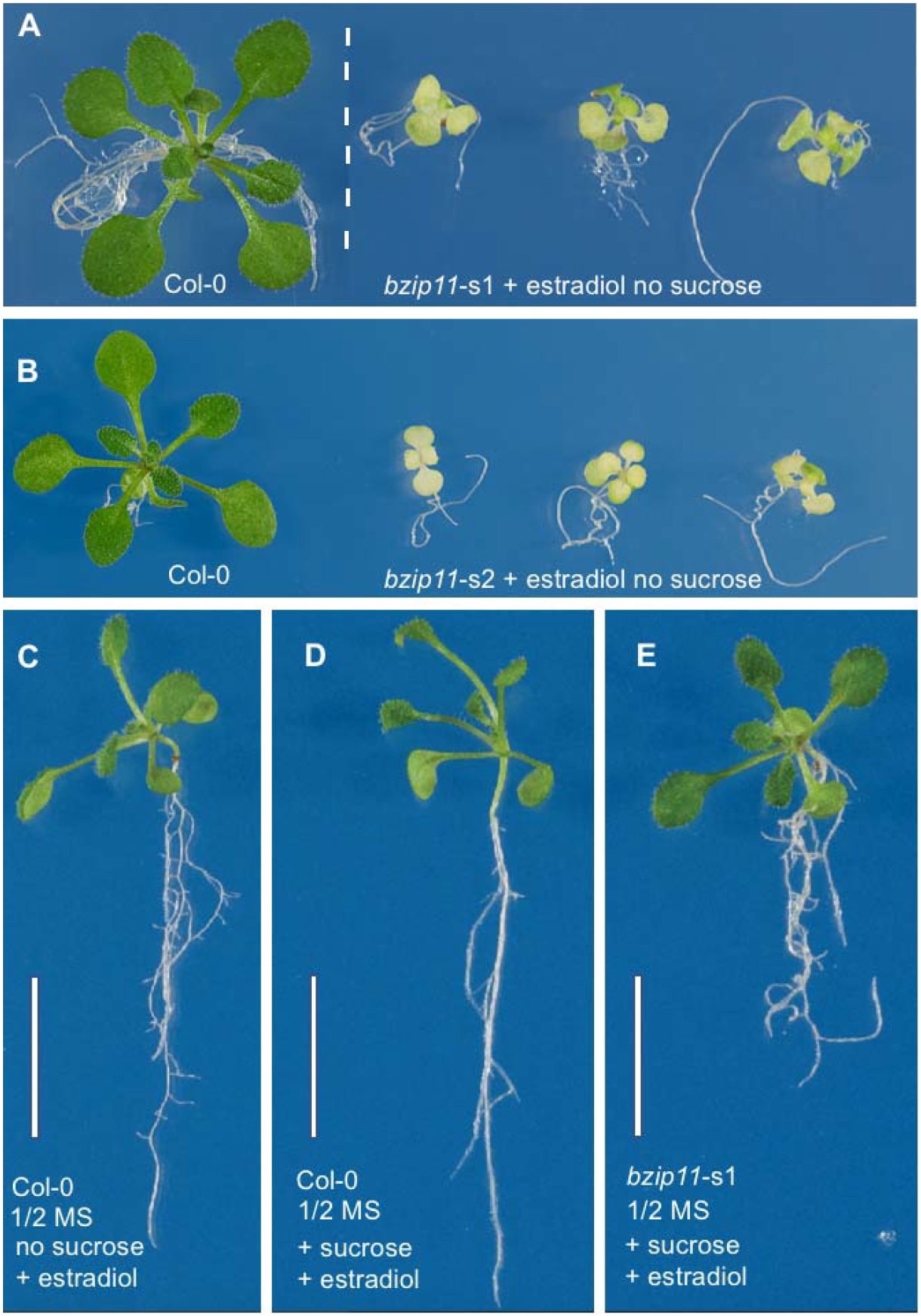
*bZIP11 amiRNA* inducible expression lines’ growth with presence or absence of sucrose and estradiol. (*A & B*) Twenty-one day old wild-type (Col-0), *bzip11*-s1 and *bzip11*-s2 representative seedlings grown on media supplemented with 50μM estradiol. (*C*) Wild-type grown on media supplemented with 50μM estradiol; (*D*) Wild-type grown on media supplemented with 50μM estradiol and 30mM sucrose; (*E*) Transgenic *bzip11*-s1 grown on media supplemented with 50μM estradiol and 30mM sucrose. Scale bar – 1cm.

To identify genes regulated by bZIP11, mRNAs differentially regulated in Col-0 relative to the *bZIP11* mutants were compared (microarray data for *bzip11*-s1 and RNAseq data for *bzip11*-s2). An unbiased gene ontology (GO) analysis was performed (*e*-Xtra 1). GO analysis of altered gene expression in *bzip11*-s1 indicates that several categories of genes connected to enzyme activity and transport are enriched such as oxidoreductase, peroxidase and transmembrane transport. Other terms that are highly enriched in the dataset are heme and tetrapyrrole binding, responses to various stresses (chemical, hypoxia; *e*-Xtra 1).

### Validation of Sequencing Experiments

A subset of the genes altered in sequencing experiments was validated by real time PCR, to confirm the downregulation patterns for *bzip11-s1* (Fig. S3). We cannot exclude that immunity may also be playing a role in the mutants as knockdown of *bZIP11* gene expression also alters a significant amount of immunity related transcripts, (*e*-Xtra 1). At the same time, the abundance of mRNAs for several transporters, including five *UmamiT*s, was reduced in the microarray analysis of *bzip11-s1* in the presence of 50 μM estradiol (Table 1). UmamiT14, UmamiT18, and UmamiT29 play critical roles in supplying amino acids to the developing seed (Ladwig et al. 2012; Müller et al. 2015). The abundance of both *UmamiT18* and *UmamiT29* mRNAs were increased in wild-type *Arabidopsis* leaves during *P. syringae* pv. *tomato* strain DC3000 induction (Kemmerling et al. 2007). The abundance of mRNAs for representatives from two types of nitrate/peptide, ammonium, copper, phosphate and zinc transporters were also reduced in both *bZIP11* amiRNA lines (Table 1). However, under the conditions tested, none of the *SWEET*s were significantly downregulated in the amiRNA lines. It is at present unclear why we did not observe significant changes in *SWEET* gene expression in these experiments. At this point, we cannot rule out the possibility that the time points chosen, or the estradiol dose used for estradiol-triggered repression of *bZIP11* expression did not capture effects on *SWEET* gene expression. These studies show that UmamiTs and a set of transporters are down-regulated in both the *bzip11*-s1 and *bzip11*-s2 mutants.

**Table 1.**
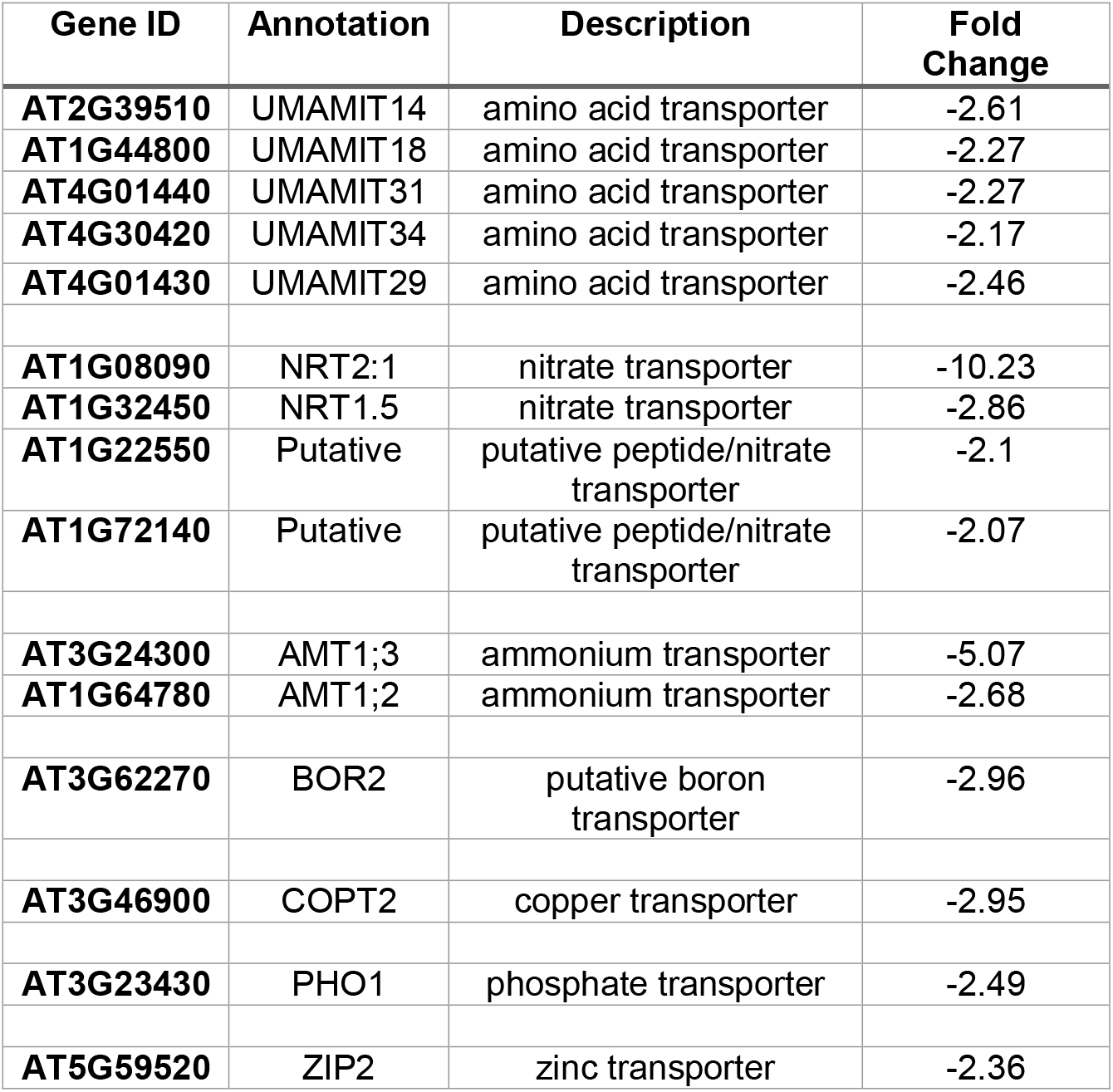
Nutrient transporters downregulated in *bzip11-*s1 knockdown transgenic plants compared to the wild-type plants. Fold change is equal to the average signal wild-type / average signal of *bZIP11-*s1. The negative sign represents a downregulation of the transporter’s expression compared to the wild-type control.

| Gene ID | Annotation | Description | Fold Change |
| --- | --- | --- | --- |
| **AT2G39510** | UMAMIT14 | amino acid transporter | -2.61 |
| **AT1G44800** | UMAMIT18 | amino acid transporter | -2.27 |
| **AT4G01440** | UMAMIT31 | amino acid transporter | -2.27 |
| **AT4G30420** | UMAMIT34 | amino acid transporter | -2.17 |
| **AT4G01430** | UMAMIT29 | amino acid transporter | -2.46 |
| **AT1G08090** | NRT2:1 | nitrate transporter | -10.23 |
| **AT1G32450** | NRT1.5 | nitrate transporter | -2.86 |
| **AT1G22550** | Putative | putative peptide/nitrate transporter | -2.1 |
| **AT1G72140** | Putative | putative peptide/nitrate transporter | -2.07 |
| **AT3G24300** | AMT1;3 | ammonium transporter | -5.07 |
| **AT1G64780** | AMT1;2 | ammonium transporter | -2.68 |
| **AT3G62270** | BOR2 | putative boron transporter | -2.96 |
| **AT3G46900** | COPT2 | copper transporter | -2.95 |
| **AT3G23430** | PHO1 | phosphate transporter | -2.49 |
| **AT5G59520** | ZIP2 | zinc transporter | -2.36 |

### *P. syringae* pv. *tomato* strain DC3000 growth is impaired in *bZIP11* mutants

To determine if silencing of *bZIP11* would affect susceptibility of *Arabidopsis* to bacteria, *bzip11*-s1, *bzip11*-s2 and Col-0 leaves were infected with a suspension of *P. syringae* pv. *tomato* strain DC3000 containing 5 μM estradiol. At three days post-infection, *bzip11*-s1 and *bzip11*-s2 leaves contained approximately 10-fold lower *P. syringae* pv. *tomato* strain DC3000 titers compared to wild-type leaves (Fig 3A). Prior to infection, leaves of *bzip11-*s1 and *bzip11-*s2 or wild-type control did not show apparent signs of chlorosis, a phenotype that could indicate stress Fig. S2B). Chlorosis was observed for mutants when grown for twenty-one days on plates with estradiol in the absence of sucrose (Fig 2A, 2B). At three days post-infection, all three adult plants showed a similar degree of pathogen-induced leaf collapse and yellowing at the leaf margins (Fig S4), typical effects observed for leaves due to the high inoculum of the pathogen. Notably, only the inducible repression of *bZIP11* expression upon infiltration with *P. syringae* pv. *tomato* strain DC3000 significantly reduced pathogen growth in leaves. Our findings indicate that bZIP11 expression in adult plants is required for maximal growth of *P. syringae* pv. *tomato* strain DC3000 in leaves. Next, we directly tested immune responses during *P. syringae* pv. *tomato* strain DC3000 infection in *bzip11*-s1 and *bzip11*-s2 by quantifying the growth of *P. syringae* pv. *tomato* strain DC3000 Δ*hrcU*, a strain that grows poorly in leaves due to the lack of a functional T3SS (26). Titers of *P. syringae* pv. *tomato* strain DC3000 Δ*hrcU* were similar in *bzip11*-s1, *bzip11*-s2, and Col-0 indicating that the *bzip11* amiRNA lines did not display enhanced resistance (Supplementary Fig. S5). Reactive oxygen species (ROS) production in leaves also appeared unaffected (Supplementary Fig. S6). The reduction in growth of *P. syringae* pv. *tomato* strain DC3000 in *bzip11*-s1 and *bzip11*-s2 compared to that in Col-0 is therefore not likely caused by an enhanced host immune response. Rather, these findings suggest that reduced pathogen growth is due to the loss of bZIP11-dependent transcription programs, which are required to promote pathogen replication.

**Figure 3.**
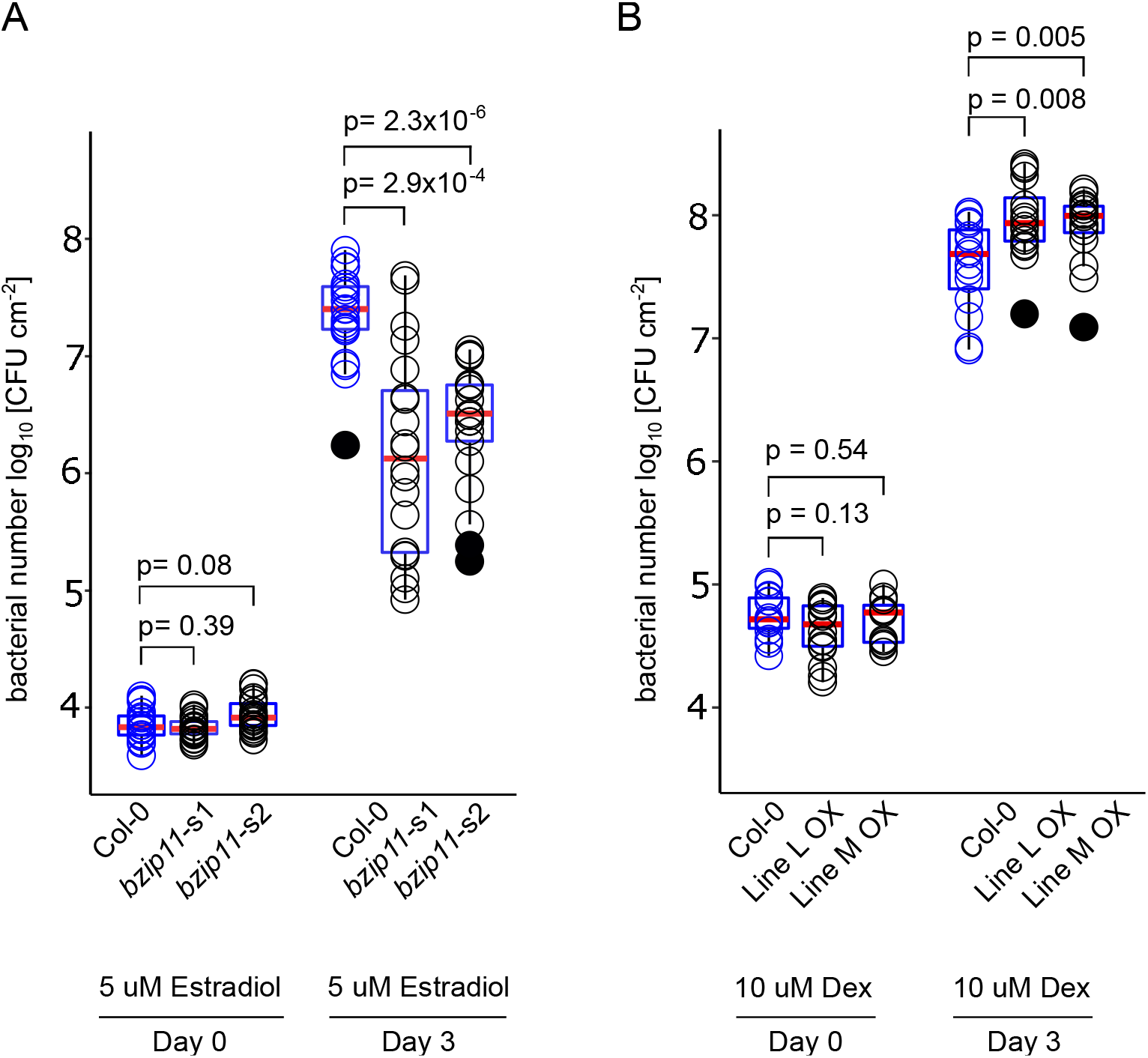
(*A*) Box plot showing levels of *P. syringae* pv. *tomato* strain DC3000 in the leaves of *Arabidopsis* wild-type Col-0 or *bZIP11* inducible amiRNA lines, *bzip11*-s1 and *bzip11*-s2. Bacterial titer at Day 0 (start of infection) and Day 3. Bacterial suspensions contained 5 μM estradiol. Data represents log10 CFU/cm^2^ from 61 individual plants. Experiments were repeated in four independent biological repeats. Blue boxes represent the 25th percentile (top) and 75 percentile (bottom) of the sample set respectively. The red line in each box is the median sample. Each circle is an independent CFU/cm^2^ count from a single leaf of one of the 61 individual plants, wild-type counts are in blue and amiRNA counts are in black. Closed circles represent sample outliers. Statistical significance by a two-way student’s t test is provided. (*B*) Box plot showing levels of *P. syringae* pv. *tomato* strain DC3000 in the leaves of *Arabidopsis* wild-type Col-0 or bZIP11 dex inducible overexpression. Bacterial titer at Day 0 (start of infection) and Day 3. Bacterial suspensions contained 10 μM dex. The data are from three independent replicates except the Day 0 for Line M is only two replicates. Blue boxes represent the 25th percentile (top) and 75 percentile (bottom) of the sample set respectively. The red line in each box is the median sample. Each circle is an independent CFU/cm^2^ count from a single leaf of one of the 47 individual plants. Statistical significance by a two-way Student’s t test is provided.

### *P. syringae* pv. *tomato* strain DC3000 growth is enhanced in *bZIP11* overexpression lines

Next we tested if overexpression of bZIP11 is sufficient to increase the growth of the pathogen. To test this, we used the previously established dex inducible Line L and Line M (Hanson et al. 2008). We infected Col-0 Line L and Line M with a suspension of *P. syringae* pv. *tomato* strain DC3000 containing 1 μM dex but observed no statistically significant change in pathogen growth across the three genotypes. However, we note that although on average there were slightly more bacteria in the overexpression lines than the Col-0 control (Figure S7). The dex concentration was next increased to 10 μM and we observed approximately a 240% and 189% increase in pathogen titer in the Line L and Line M lines, respectively, compared to the Col-0 control at three days post-infection (Figure 3B). The Line L had on average a 27% higher titer of bacteria than the Line M. This finding is consistent with the previous observation that the Line L displayed stronger expression of the transgene than Line M, although this difference in bacterial titer was not statistically significant (Hanson et al. 2008). The dose dependent increase in *P. syringae* pv. *tomato* strain DC3000 growth in the dex inducible *bZIP11* lines compared to the Col-0 control further suggests that overexpression of bZIP11 is sufficient to increase pathogen growth. Thus, we conclude that bZIP11 is a host susceptibility factor that impacts the outcome of bacterial infection in *Arabidopsis*.

## Discussion

The strategies that allow pathogens to acquire nutrients from the host to sustain invasion and reproduction in host plants are still largely unknown. The understanding of the bacterial gene-for-gene regulatory system that allows *Xanthomonas* species to infect a variety of plant hosts is now a basis for engineering resistance in rice and specialized diagnostic toolkits for *Xanthomonas oryzae* pv. *oyzae* infection (Chen et al. 2010; Eom et al. 2019; Oliva et al. 2019). As there are hundreds of economically important plant pathogens and only a few are known to possess TAL like effectors, the *Xanthomonas* TAL effector – host *SWEET* gene for gene susceptibility mechanism is more likely an exception rather than a general rule. Hijacking the transcriptional programs of secretory cell types may represent an alternate mechanism to access a broad spectrum of plant nutrients beyond soluble carbohydrates.

Overexpression of bZIP11 or bZIP11 homologs led to a marked increase in sugars and amino acids in several different plants systems, indicating they influence downstream nutrient pools (Hanson et al. 2008; Ma et al. 2011; Thalor et al. 2012). The microarray and RNAseq analyses of wild-type and *bzip11* mutants revealed that bZIP11 is required for the expression of multiple nutrient transporter genes, as well as genes involved in metabolic processes and growth (Table 1, Table S4). Similarly, previous studies reported that overexpression of bZIP11 results in an overall increase in sugars and multiple amino acids relative to wild-type plants (Hanson et al. 2008). Notably, *Pseudomonas syringae* pv. *tomato* strain DC3000 growth was significantly increased in leaves overexpressing bZIP11 (i.e. bZIP11 dex inducible lines treated with 10 μM dex) compared to WT Col-0 lines. These data indicate that activation of bZIP11-dependent nutrient pathways increases susceptibility to *Pseudomonas syringae* pv. *tomato* strain DC3000. These data are consistent with our model that bacterial modulation of host *bZIP11* mRNA expression results in an increase in available nutrient pools and the expression of a wide range of transporters.

Interestingly, alteration of host nutrient transporter expression seems to be crucial in several host-pathogen systems. The *Arabidopsis* hexose/H^+^ symporter STP13 was found to compete for sugars in the apoplasm during *P. syringae* pv. *tomato* strain DC3000 infection as a key part of the host’s defense response, consistent with the idea that competition over nutrient availability is a crucial aspect of the host-pathogen relationship (Yamada et al. 2016). Additionally, mutations in the wheat *STP13* homolog confer broad-spectrum resistance against fungal rusts (Moore et al. 2015). Recently the unmutated *TaSTP13* gene in wheat was shown to confer susceptibility to stripe rust, potentially by increasing apoplasmic hexose availability (Huai et al. 2020). In rice, *OsSWEET13* and *OsSWEET15* were reported to be directly regulated by OsbZIP73 and induced during salt and drought stress (Mathan et al. 2020). In our sequencing experiments however, none of the *SWEET*s were significantly downregulated in the two amiRNA lines. Interestingly, a previous study reported that the ectopic overexpression of bZIP11 in the Line M and Line L seedlings resulted in the downregulation of *SWEET1* and *SWEET12* mRNA levels when the seedlings were treated with 10 μM dex induction for 2 hours (Hanson et al. 2008). Additional work may help distinguish the differences between rice and *Arabidopsis* with regards to the bZIPs and *SWEET* regulation. The modulation of host nutrient transporters and the accompanying changes in metabolite availability have also been proposed to play important roles in the host immune response (Herbers et al. 1996; Kocal et al. 2008; Sonnewald et al. 2012). Interestingly, a pepper bZIP protein, CabZIP2, is required for full resistance in pepper against *X. euvesicatoria*, and transgenic overexpression of CabZIP2 in *Arabidopsis* increased resistance to *P. syringae* pv. *tomato* strain DC3000 (Lim et al. 2015). Thus, it is possible that bZIP proteins may play roles in either resistance or susceptibility in different pathosystems.

The restriction of pathogen growth in the *bZIP11* mutants does not appear to be an enhanced resistance response. Testing a direct link between pathogen growth and the host’s nutritional status remains challenging because we predict that pathogen induction of *bZIP11* expression alters not only nutrient transporter expression but also the metabolic state of the infected cells. Overexpression of candidate nutrient transporters alone is thus not likely to be sufficient for rescuing the *bzip11* mutant phenotype in pathogen-infected leaves. Moreover, due to the induction of many transporters, overexpression of a single gene will not likely lead to complementation of the phenotype.

In conclusion, our findings suggest that bZIP11 functions as a susceptibility factor during *P. syringae* pv. *tomato* strain DC3000 infection in *Arabidopsis*. Reduced *bZIP11* expression limits *P. syringae* pv. *tomato* strain DC3000 growth, while increased *bZIP11* expression increases growth. These data indicate that bZIP11-regulated metabolic networks may provide nutrients to fuel pathogen reproduction at the local infection site or to regulate host defense responses. Further insight into the role of plant secretory programs in plant disease susceptibility and how they may be co-opted by pathogens may offer new strategies to engineer pathogen resistance in crop plants.

## Materials and Methods

### Bioinformatics

Comparison of 76 *Arabidopsis* seed developmental transcription factors and 74 *Arabidopsis* transcription factors induced in leaves by *P. syringae* pv. *tomato* strain DC3000 was completed in a Microsoft Excel to identify candidate *bZIP11*. Analysis of *SWEET* and *UmamiT* promoters was completed using Pscan (Zambelli et al. 2009).

### DNA constructs

The *bZIP11* artificial microRNA (amiRNA) sequence was generated using the Weigelworld online web MicroRNA designer (http://wmd3.weigelworld.org/cgi-bin/webapp.cgi). The amiRNA precursor was produced in pRS300 using published protocols (Schwab et al. 2006). The modified amiRNA precursor was inserted into pDONOR221 (Thermo). The estradiol-inducible promoter G1090 and the *NOS* terminator were amplified and cloned into pDONORp4p1r and pDONORp2rp3. PCR amplification was performed with Phusion High-Fidelity DNA Polymerase (Thermo) and confirmed by sequencing and integrated into destination vector pB7m34GW-R4R3 by multisite gateway technology (Invitrogen). The final recombinant destination vector was confirmed by restriction profile and mobilized into *Agrobacterium tumefaciens* C58C1 by electroporation. The plasmid DNA was isolated from agrobacteria and confirmed by restriction analysis before transformation.

### Plant material and transformation

*Arabidopsis thaliana* (Col-0) plants were grown on half-strength Murashige-Skoog (MS) media (Duchefa, Haarlem, Netherlands) supplemented with 0.8% agar, pH 5.8. The seeds were sterilized by chlorine gas and were stratified for 3-4 days at 4°C in the dark. Ten-day-old seedlings were transferred to soil (50% compost, 50% vermiculite) and cultivated under fluorescent light (16-h light, 120 μE m^−2^ s^−1^) until flowering. The floral dip transformation method was used to stably transform *Arabidopsis* plants (Clough and Bent 1998). Transgenic plants were selected on half strength MS media supplemented with 0.8% plant agar (Duchefa) and the appropriate BASTA (Duchefa) before transfer to soil. T2 selfed progeny were tested for segregation of the selectable marker on solid half strength MS media. Only seedlings of lines showing a 3:1 segregation were considered for further analysis. Homozygous *bzip11*-s1 and *bzip11*-s2 lines were generated by selfing and used for further experimentation. Seedling growth phenotypes were monitored on half strength MS medium with 50 μM estradiol ± 30 mM sucrose (16-h light, 120 μE m^−2^ s^−1^).

### RNA and real-time PCR analyses

Plant material was frozen in liquid nitrogen and ground using glass beads in a Micro-Dismembrator. Total RNA was isolated and purified using the RNeasy kit (Qiagen) and quantified by NanoDrop. 1 μg of DNase treated RNA was used for cDNA synthesis by oligo-dT primers and MLV reverse transcriptase (Promega) enzyme. The Applied Biosystems with ABIPRISM® 7900 was used for gene quantification using SYBR® Green and used a standard PCR program for amplification (50°C for 2 min, 95°C for 10 min, 40 cycles of 95°C for 15 s, and 60°C for 1 min). Amplicon dissociation curves were recorded after 40 cycles by heating from 60 to 95°C with a ramp speed of 1.9°C min^−1^. Target genes transcript levels were calculated relative to the reference gene At1g13320 (Czechowski et al. 2005). RNA was isolated from *P. syringae* pv. *tomato* strain DC3000 infected leaves using Sigma Aldrich’s Plant Total RNA isolation kit and protocol. For each sample, 1 μg of total RNA was reverse transcribed using the Quantitech Reverse Transcription kit and protocol. Quantitative PCR analysis was performed using the Syber Green real time PCR kit. Primers were designed using the Primer3 software (Rozen and Skaletsky 2000). All primer pairs used are present in Supplementary Table S2.

### Microarray analyses

Total RNA was isolated from frozen tissues of seven day old seedlings treated with 50 μM estradiol using a Sigma Aldrich-Spectrum Plant Total RNA Kit and quantified by Nanodrop ND-1000 Spectrophotometer analysis. Three biological replicates were used for microarray analysis. The microarray experiments were performed by ServiceXS (www.servicexs.com) using AffymetrixATH1 GeneChips. Quality control of scanned arrays was analyzed using R language, Bioconductor (https://www.bioconductor.org/) and Affymetrix Expression Console Software (http://www.affymetrix.com/). Bioconductor was used for Robust Multi-array Average (RMA) normalization of raw data at gene level to obtain signal intensity values. Limma package was used for comparing treatments and differential expression analysis (Smyth 2005). For functional analysis, g:profiler was used (https://biit.cs.ut.ee/gprofiler/gost). Raw data can be found at https://www.ncbi.nlm.nih.gov/geo/ under accession nr GSE139821

### RNAseq analyses

Three replicates of the RNA samples from seven day old seedlings treated with 50 μM estradiol were used for library preparation (TruSeq RNA Sample Prep Kit v2) and sequenced using Illumina HiSeq2500, single read, 1×50bp. Library preparation and sequencing was performed by Macrogen Europe BV (Amsterdam, Netherlands). Raw sequencing reads were aligned to the *Arabidopsis* genome (TAIR10) using TopHat v2.0.13 (Trapnell et al. 2009) with the parameter settings: ‘bowtie1’, ‘no-novel-juncs’, ‘p 6’, ‘G’, ‘min-intron-length 40’, ‘max-intron-length 2000’. On average 97.0% (94.0 – 98.1%) of the RAW reads could be aligned to the genome per biological replicate. This represents an average of 51.1 (40.7 – 73.3) million mapped reads. Aligned reads were summarized over annotated gene models using HTSeq-count v0.6.1 (Anders and Huber 2010) with settings: ‘-stranded no’, ‘-i gene_id’. From the TAIR10 GTF file all ORFs (open reading frames) of which the annotation starts with ‘CPuORF’ were manually removed prior summarization to avoid misalignment of polycistronic mRNAs. Normalized counts were imported to LIMMA (Ritchie et al. 2015) for statistical determination of differential gene expression. Genes for the normalized count that changed more than twofold with an associated *P*-value of less than 0.05 (Bonferroni Holm test corrected) were considered to be differentially expressed. For functional analysis, g:profiler was used (https://biit.cs.ut.ee/gprofiler/gost). Raw data can be found at https://www.ncbi.nlm.nih.gov/geo/ under accession nr GSE135593.

### Bacterial infection assays

For infection assays with *P. syringae* pv. *tomato* strain DC3000, seeds were vernalized for two days at 4°C. Plants were grown for 6.5 weeks in a Percival growth chamber, model I22NLC8, set at a 10-h light (175 μE m^−2^ s^−1^ at 22°C), 14-h dark cycle at 80% humidity and watered once a week. For the dex inducible bZIP11 overexpression line studies, a Percival growth chamber, CU36L4 was used with the same settings. Plants were fertilized with Peters Excel water soluble 15-5-15 (Scotts-Sierra Horticultural Products Company, Marysville, Ohio) at week 2, then every other week until 6.5 weeks of age. Plants were selected such that all Col-0 wild-type, *bzip11-*s1 and *bzip11-*s2 plants within each genotype were of similar size, to control for variation in plant size resulting from leaky *bZIP11* expression among individual plants. The Col-0 control and *bzip11-*s1 individual plants chosen for infection were of comparable size at 6.5 weeks of growth while the bzip11-s2 plants were slightly smaller. Plants were inoculated with a 2×10^6^ CFU/ml suspension of *P. syringae* pv. *tomato* DC3000 in 1 mM MgCl_2_ and 5 μM estradiol, 1 μM, or 10 μM dexamethasone. No drug was used for real time PCR analysis of infected tissue. The leaf samples for the qPCR analysis were collected from the same inoculated plants at three time points. A minimum of three individual plants were used for each indepdenent biological replicate. Each time point was obtained by harvesting at least three individual leaves from at least three individauls plants and pooling them for a single sample for that specific time point). The mock was infiltration with 1 mM MgCl_2_. Infected plants were placed in a covered flat with 200 ml of standing water and incubated for 3 days for the growth assay. Cork borers (4.88 mm diameter) were used to isolate four leaf discs from each leaf. The leaf discs were ground and suspended in 1 mM MgCl_2_ for manual CFU counting. For the qPCR analysis in infected leaves

### Oxidative burst assays

Three soil grown, 4-week-old plants were selected for each line. Two leaves per plant were harvested from the youngest fully expanded leaves. Two leaf discs (5-mm diameter) taken from each harvested leaf (n = 12) and incubated in water in a 96-well plate (one leaf disc per well) for 24-h. To measure ROS, leaf discs were treated with flg22 (100 nM) in a 10 μg/mL horseradish peroxidase and 100 μM Luminol (Sigma) solution, and then immediately subjected to luminescence measurement with a 1420 Multilabel Counter (PerkinElmer). Relative luminescence units (RLU) are reported. Assay was repeated three times independently with similar results.

## Supporting information

Raw Data location

e-Xtra file 1

## Acknowledgements

MJP was additionally supported by a Stanford Graduate Fellowship (SGF) fellowship. We thank Dr. Alexander Jones (University of Cambridge), Dr. Li-Qing Chen (University of Illinois Urbana-Champaign), and Dr. Davide Sosso for many useful discussions.

MJP, JS, MBM, SS, JH, WBF designed research plans. MJ, MBM, SS, JH, WBF supervised experimental work. MJP, JS, JGK, MT performed experiments. MJP, JS, MBM and WBF completed the writing with contributions from all other authors.

This work was supported by the National Science Foundation research grants IOS-1258103, the Office of Basic Energy Sciences of the US Department of Energy grant number DE-FG02-04ER15542, Deutsche Forschungsgemeinschaft (DFG, German Research Foundation) under Germany′s Excellence Strategy – EXC-2048/1 – project ID 390686111 and the Alexander von Humboldt Professorship to WBF, the National Science Foundation grant (IOS-1555957) to MBM, and the Netherlands Organization for Scientific Research (NWO; grant #854.10.011) to SS.

## SUPPLEMENTARY MATERIAL

**Supplementary Fig. S1.** Analysis of *bZIP11* expression in wild-type Col-0 during infection by *P. syringae* pv. *tomato* strain DC3000.

**Supplementary Fig. S2.** Levels of *bZIP11* repression in amiRNA lines compared to wild-type and growth of amiRNA lines and wild-type in soil.

**Supplementary Fig. S3.** Validation by qPCR of microarray expression data.

**Supplementary Fig. S4.** Representative images of Col-0, *bzip11*-s1, *bzip11*-s2 leaves inoculated with *P. syringae* pv. *tomato* strain DC3000 and 5 μM estradiol concentration in the inoculum at three days post-infection. Scale bar is 1 cm. Results were similar for four independent biological replicates.

**Supplementary Fig. S5.** Growth assay of *P. syringae* pv. *tomato* strain DC3000 Δ*hrcU* in amiRNA lines with and without estradiol.

**Supplementary Fig. S6.** ROS production assay in amiRNA lines and wild-type control plants with and without estradiol.

**Supplementary Fig. S7.** Growth assay of *P. syringae* pv. *tomato* strain DC3000 in Col-0 control and dex inducible overexpression lines, Line L and Line M at 1 μM dex concentration. Data are from three independent biological replicates. Data represents log10 CFU/cm^2^ from 35 individual plants, wild-type counts are in blue and overexpression counts are in black. Bacterial titer at Day 0 (start of infection) and Day 3. Bacterial suspensions contained 1 μM dex.

**Supplementary Table S1.** Summary of expression data from two studies to identify four initial candidate transcription factor genes.

**Supplementary Table S2.** Summary of cis regulatory elements (CREs) scan of promoters of selected *UmamiT* and *SWEET* genes.

**Supplementary Table S3.** qPCR primers used in this study.

***e*-Xtra 1.** Full analysis of microarray and RNAseq expression data for mutant inducible knockdown lines when treated with 50 uM estradiol and full GO analysis of the expression data with mutant lines.

---

**Supplementary Table S1.**
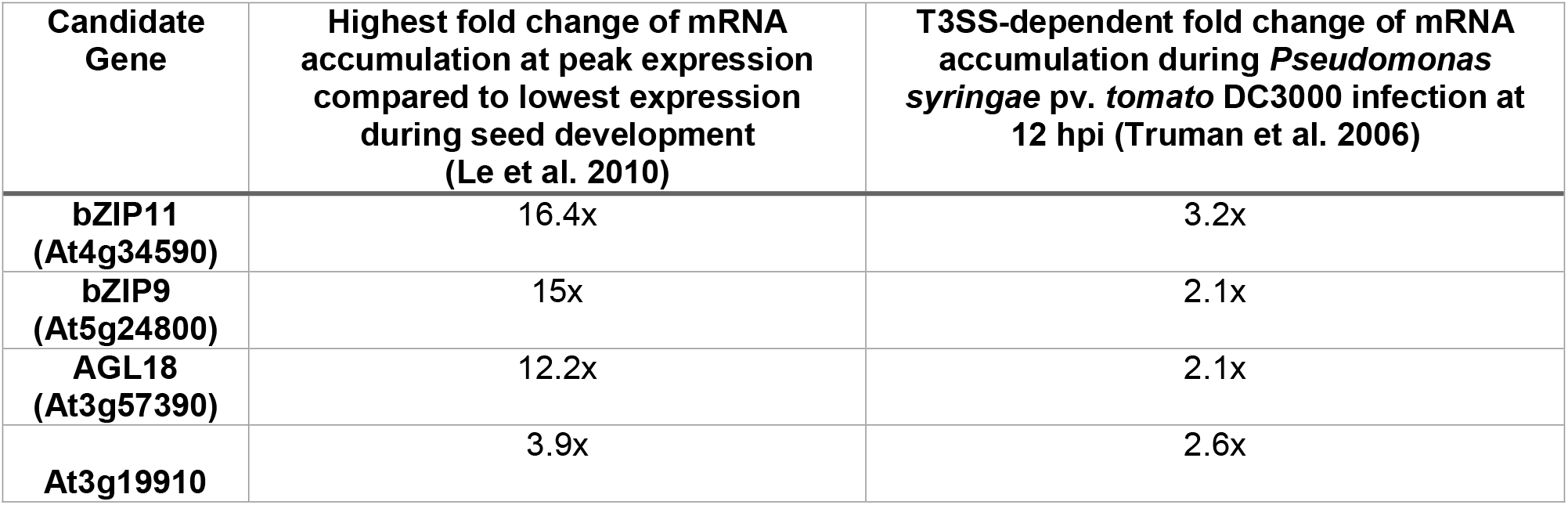
Summary of expression data from two studies to identify four initial candidate transcription factor genes.

**Supplementary Table S2.** Scan of promoters of selected *UmamiT* and *SWEET* genes for cis regulatory elements (CREs). P-values of the *UmamiT* and *SWEET* promoters are shown in comparison to two positive control promoters that have experimentally been confirmed as bZIP11 targets – ASN1 and ProDH2 – using Pscan (Hanson et al. 2008). The P-value represents how many times the given CRE would have a higher matching score in the whole genome promoter set compared to the promoter of interest (Zambelli et al. 2009).

| Gene Name | ID | bZIP<br>CREB/G-box-like<br>subclass P<br>value | bHLH(zip)<br>class P value | Published<br>Role | Literature<br>Citation |
| --- | --- | --- | --- | --- | --- |
| ***ASN1*** | At3g47340 | 0.59 | 0.03 | Glutamine<br>dependent<br>asparagine<br>synthase | Hanson<br>2008 |
| ***ProDH2*** | At5g38710 | 0.04 | 0.04 | Proline de<br>hydrogenase |  |
| ***UmamiT18*** | At1g44800 | 0.59 | 0.03 | amino acid<br>transport to<br>siliques | Ladwig<br>2012 |
| ***UmamiT20*** | At4g08290 | 0.04 | 0.02 | unknown | - |
| ***UmamiT29*** | At4g01430 | 0.92 | 0.53 | amino acid<br>transport to<br>developing<br>seed | Müller<br>2015 |
| ***SWEET4*** | At3g28007 | 0.04 | 0.12 | sugar<br>transport to<br>axial sinks | Liu 2016 |
| ***SWEET7*** | At4g10850 | 0.04 | 0.41 | unknown | - |
| ***SWEET15*** | At5g13170 | 0.38 | 0.06 | sucrose<br>transport to<br>developing<br>seed | Chen 2015 |

**Supplementary Table S3.**
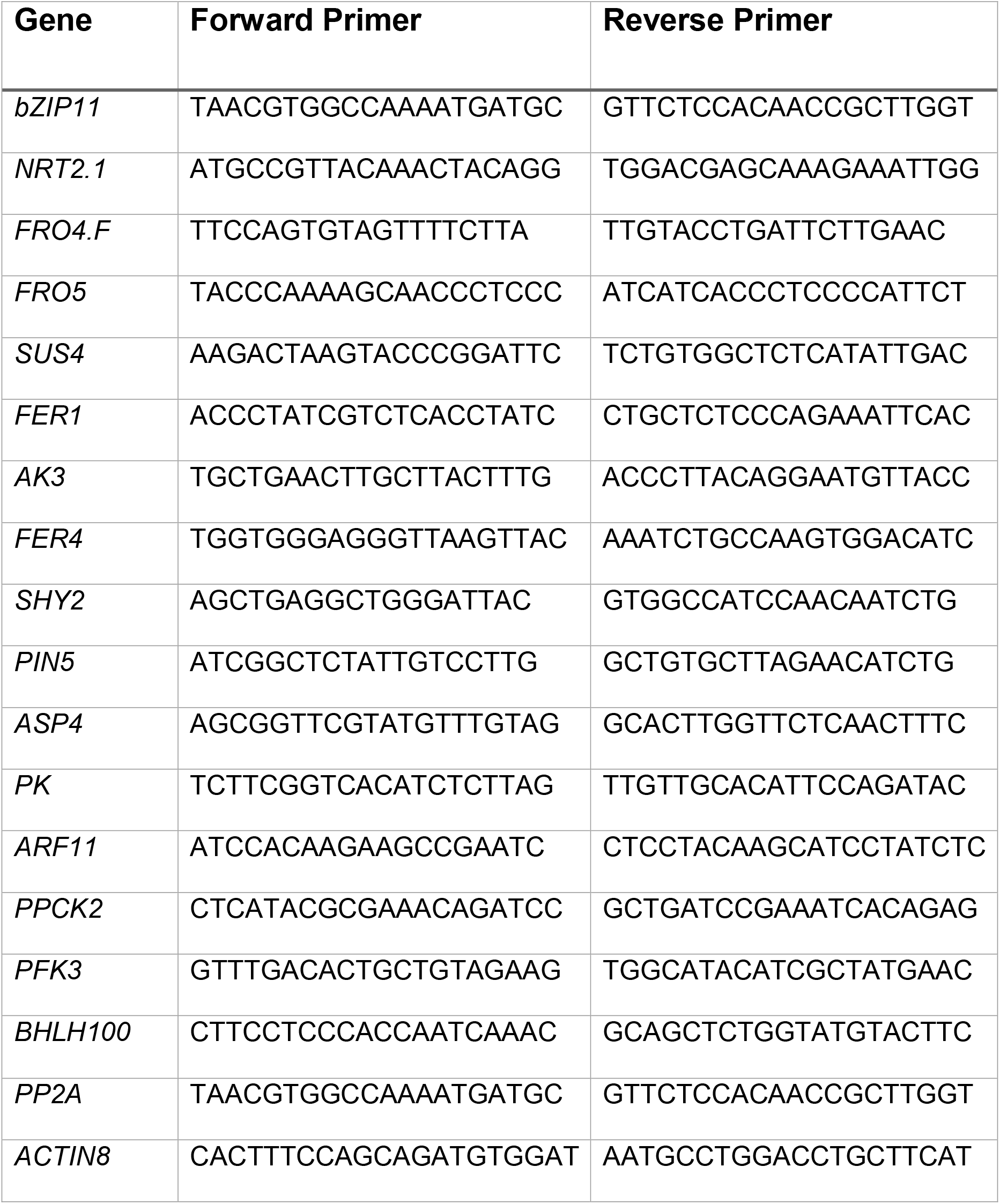
qPCR primers used in this study.

**Supplementary Figure 1.**
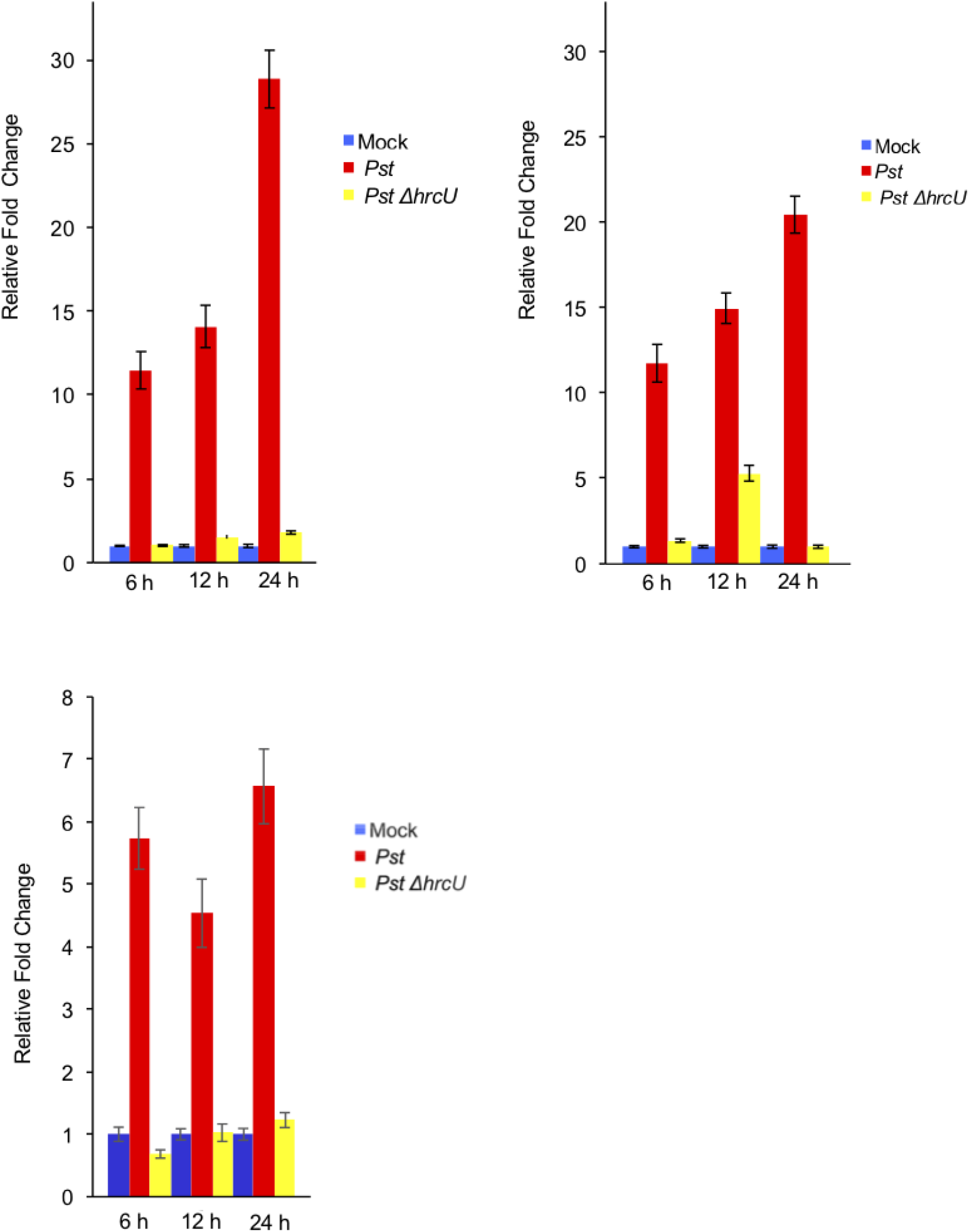
Induction of *bZIP11* expression in wild-type plants during infection by wild-type *P. syringae* pv. *tomato* strain DC3000 or the T3SS mutant Δ*hrcU* strain compared to control (mock). *bZIP11* mRNA levels were quantified by quantitative PCR (qPCR). Relative fold change (mean ± SD, n = 3 individual plants for each strain) was determined against the mean of the mock samples at each time point. All three independent biological replicates are shown.

**Supplementary Figure 2.**
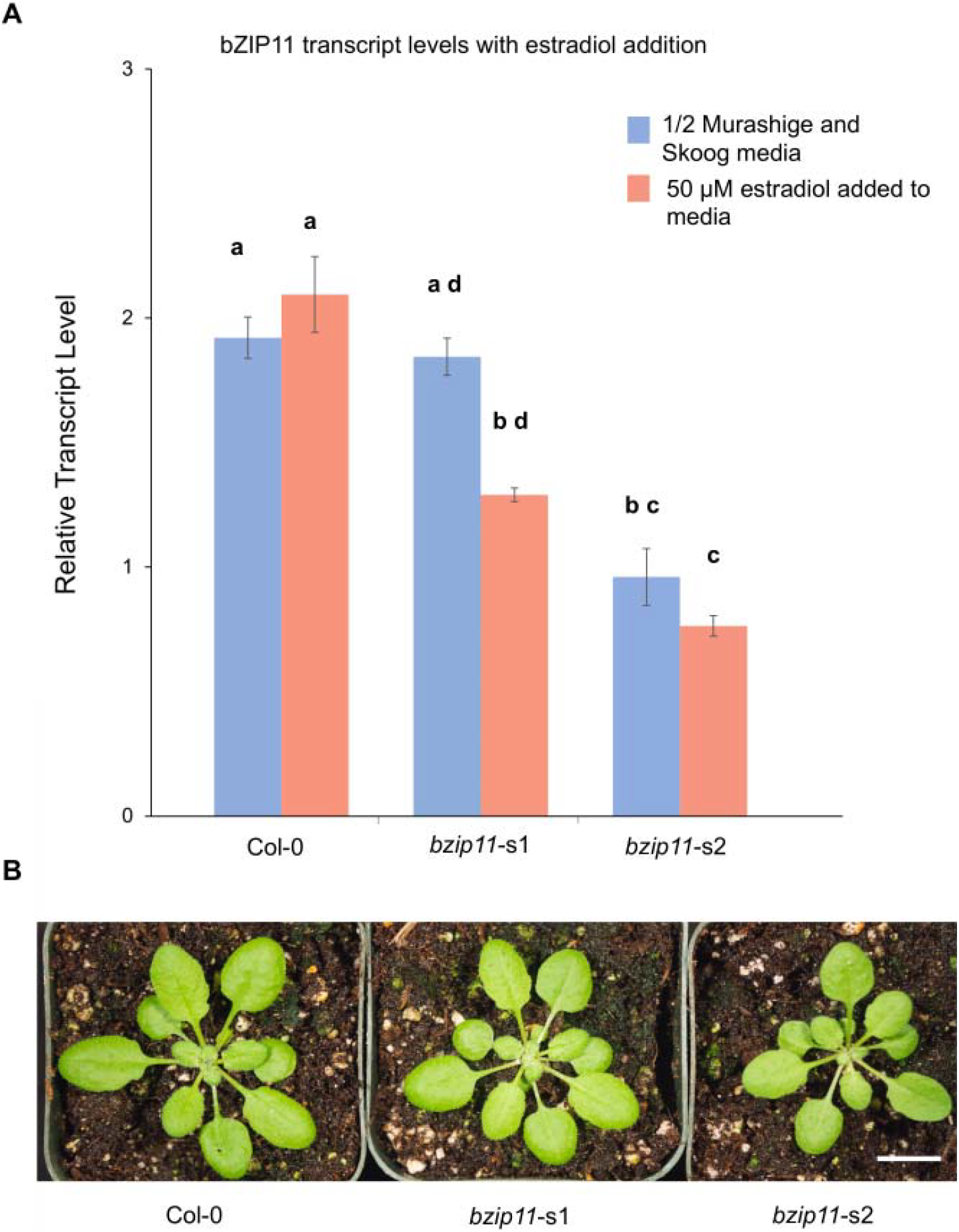
Col-0, *bzip11*-s1, *bzip11*-s2 plants were assayed for amiRNA induction and growth. *(A) bZIP11* mRNA levels with and without 50 μM estradiol treatment from seedlings. Letters indicate significant difference analyzed by one-way ANOVA followed by post-hoc Bonferroni Holm test. The reference gene use was *PP2A*. Data shown is mean ± SD, n =10. Results were repeated with three independent replicates. (*B*) Growth of Col-0, *bzip11*-s1, and *bzip11*-s2 plants without estradiol in soil. These plants are representative images of the plants used for infection at this age but these 3 individual plants were not infected. Plants are four weeks old. Scale bar – 1cm.

**Supplementary Figure 3.**
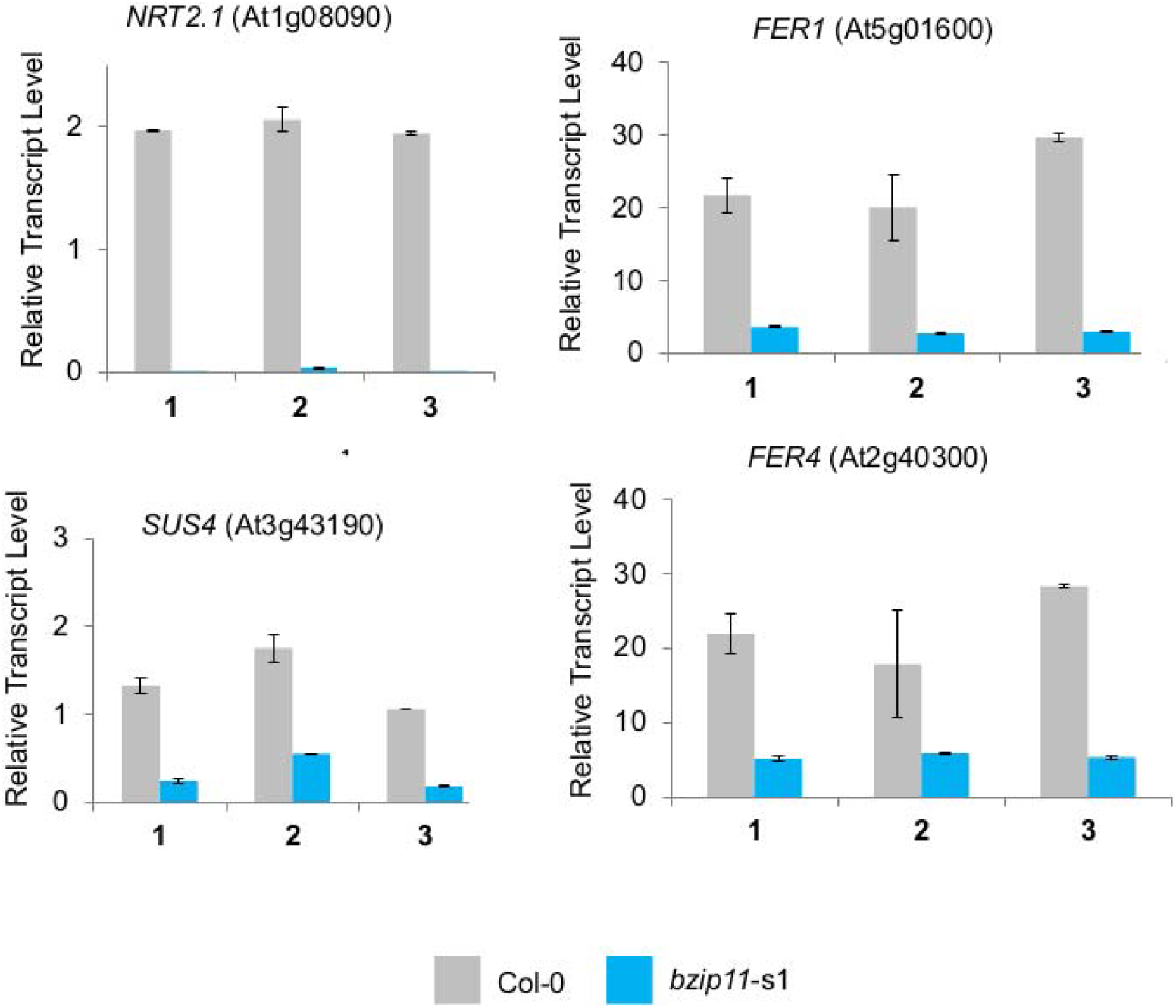
Validation of *bzip11-s1* microarray data performed by qPCR of select genes. Col-0 and *bzip11-s1* plants were treated with 50μM estradiol. Each gene was assayed independently in triplicate (1-3). Relative transcript levels (mean ± SD) are shown. N=10 per independent repeat.

**Supplementary Figure 4.**
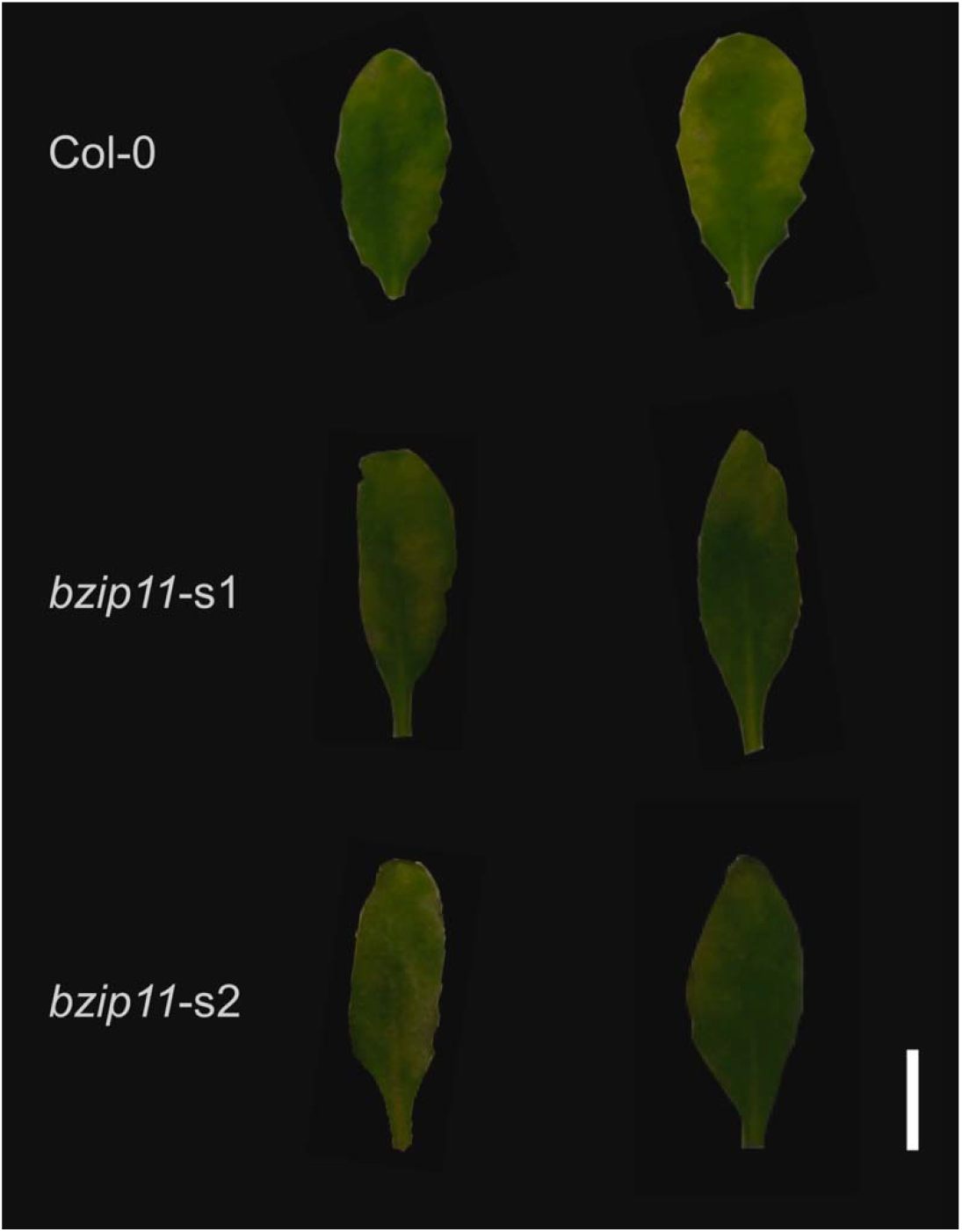
Representative images of Col-0, *bzip11*-s1, *bzip11*-s2 leaves inoculated with *P. syringae* pv. *tomato* strain DC3000 with 5 uM estradiol concentration in the inoculum at three days post-infection. Scale bar is 1 cm. Results were similar for four independent biological replicates.

**Supplementary Figure 5.**
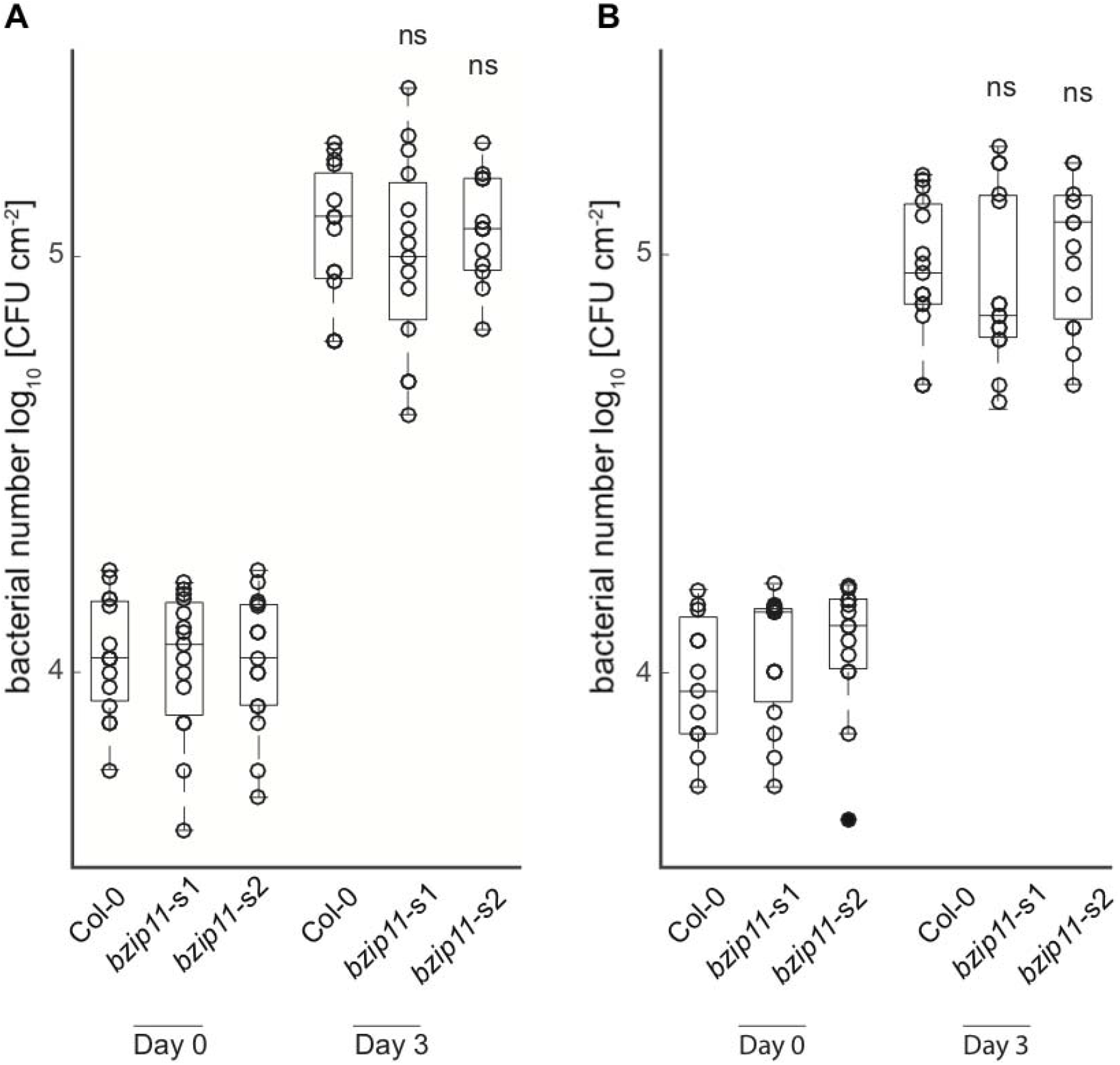
Growth assay of *P. syringae* pv. *tomato* strain DC3000 Δ*hrcU* in amiRNA lines with and without estradiol.

**Supplementary Figure 6A and 6B.**
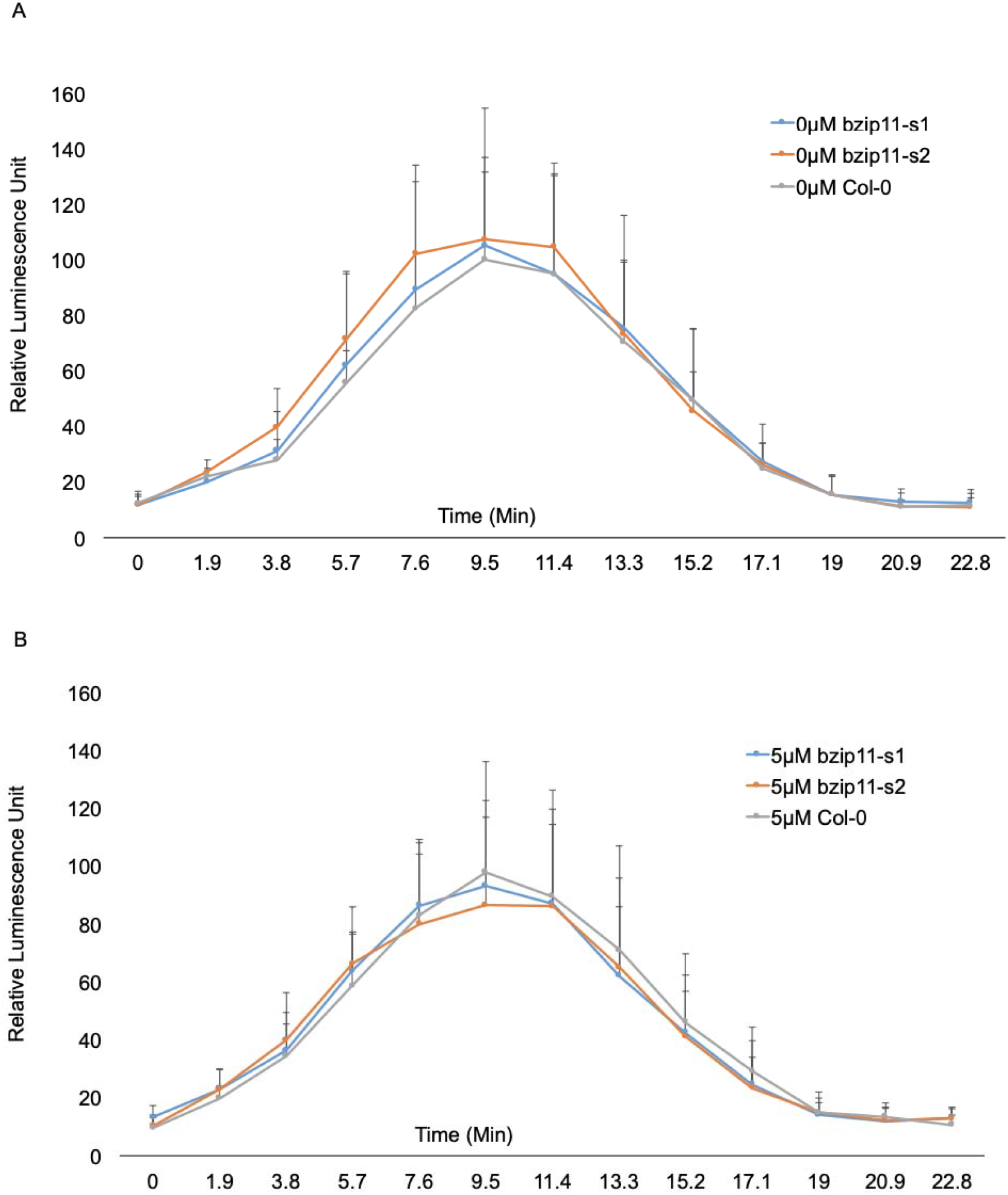
Flg22 induced ROS production in the Col-0, *bzip11*-s1, and *bzip11*-s2 lines. (A) Flg22 treatment without estradiol, (B) with 5 uM estradiol. Time course of ROS production in leaf discs treated with flg22 was measured in relative luminescence units. Error bars represent mean ± SD, n =12 leaf discs for each individual plant line. Results were similar in three separate biological repeats for each condition.

**Supplementary Figure 7.**
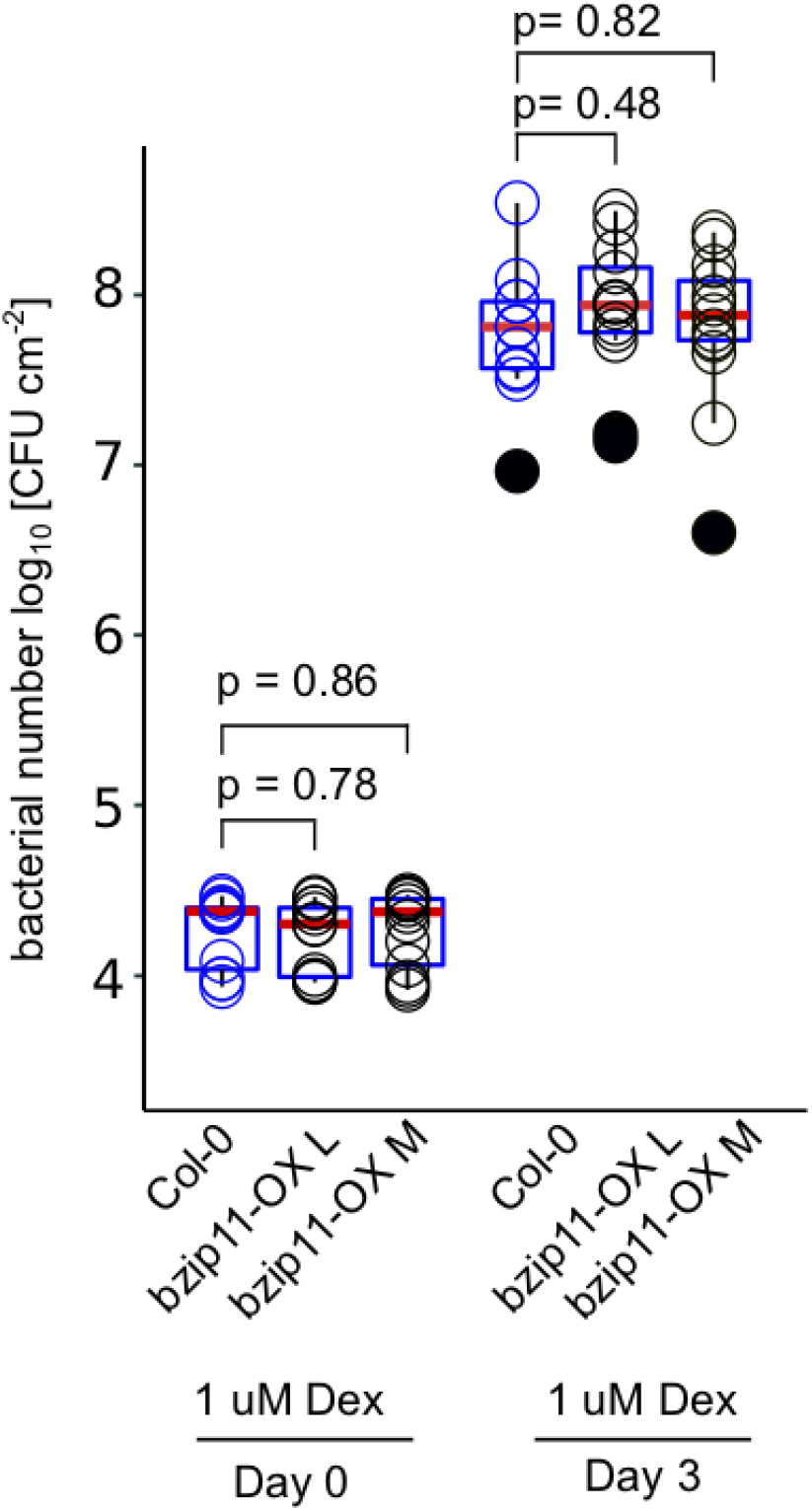
Growth assay of *P. syringae* pv. *tomato* strain DC3000 in Col-0 control and dex inducible overexpression lines, Line L and Line M at 1 uM dex concentration. Data are from three independent biological replicates. Data represents log10 CFU/cm^2^ from 35 individual plants, wild-type counts are in blue and overexpression counts are in black. Bacterial titer at Day 0 (start of infection) and Day 3. Bacterial suspensions contained 1 μM dex.

