## Supplementary material for "*Arabidopsis* bZIP11 is a susceptibility factor during *Pseudomonas syringae* infection": Raw Data location

- “e-Xtra” .xlsx file including all micro-array and RNA-seq data and a revised, unbiased GO analysis. Raw microarray data can be found at https://www.ncbi.nlm.nih.gov/geo/ under accession nr GSE139821 (enter token afihsaumjtyllyt into the box for access). The raw RNAseq raw can be accessed at the https://www.ncbi.nlm.nih.gov/geo/ under accession nr GSE135593 (enter token sbozuwoqpvyfhel into the box for access).
